# The behavioural responses of bumblebees *Bombus terrestris* in response to simulated rain

**DOI:** 10.1101/2023.12.04.569872

**Authors:** Laura A. Reeves, Ellie M. Jarvis, David A. Lawson, Sean A. Rands

**Author notes:** LAR and EMJ contributed equally, and should be treated as joint first-authors.

## Abstract

Bumblebee activity typically decreases during rainfall, putting them under the threat of the increased frequency of precipitation due to climate change. A novel rain machine was used within a flight arena to observe the behavioural responses of bumblebees (*Bombus terrestris*) to simulated rain at both a colony and individual level. During rainfall, a greater proportion of workers left the arena than entered, the opposite of which was seen during dry periods, implying that they compensate for their lack of activity when conditions improve. The proportion of workers flying and foraging decreased while resting increased in rain. This pattern reversed during dry periods, providing further evidence for compensatory activity. The increase in resting behaviour during rain is thought to evade the high energetic costs of flying whilst wet without unnecessarily returning to the nest. This effect was not repeated in individual time budgets, measured with lone workers, suggesting that the presence of conspecifics accelerates the decision of their behavioural response, perhaps via local enhancement. Bumblebees likely use social cues to strategise their energetic expenditure during precipitation, allowing them to compensate for the reduced foraging activity during rainfall when conditions improve.

## INTRODUCTION

During the next century, global temperatures are estimated to rise by several degrees [1], and multiple climate change scenarios predict that this will be accompanied by higher levels of rainfall, both on average and during extreme weather events [2,3]. This rainfall will not be evenly distributed globally, with some areas receiving more precipitation than others [4,5]. Altered levels of rainfall could disrupt a large number of processes within ecosystems such as pollination [6,7]. Rainfall may disrupt plant-pollinator signalling by acting as environmental noise: floral volatiles can be easily removed or diluted by rain and raindrops can interfere with pollinator’s sensory systems, making it difficult for pollinators to receive signals used in advertising floral rewards [7]. Floral rewards such as pollen and nectar can be diluted or degraded by rainfall, reducing the likelihood of pollinator visitation due to decreased reward quality [7]. Rain may also directly affect pollinators, *via* raindrop impacts, or by altering thermoregulation and increasing body mass, raising the amount of drag and energetic costs during flight and take-off [8].

There is a large gap in the scientific literature concerning the effects of rainfall on pollination or insect behaviour [7,9,10], potentially because behaviour during rainfall is tricky to study in the field, and technically awkward to simulate in the lab. A few studies have focussed on bee behaviour, and have found a negative correlation with worker activity [11–13], even as far as reducing the trophallactic activity between nurses and larvae inside the nest [14]. Honey bees (*Apis mellifera*) increase their foraging effort before and after rainfall to compensate for the reduction in activity [15–17], suggesting that this is less energetically costly than continuing to forage during rainfall.

In this paper, we consider how we could start systematically investigating bee behaviour, by demonstrating that rainfall simulation can be integrated into some standard bumblebee experimental techniques. Rainfall simulators are a valuable tool for exploring how rain interacts with the environment, and many different designs have been described within the hydrology and soil erosion literature [18–21], with designs either focussed on using gravity to create drips through thin tubes, or forcing pumped water through nozzles. Rain simulation has been used to explore the water-living larvae of various species of mosquito may be flushed out of their nursery pool in heavy rain [22–24], or the removal from leaves of protective structures produced by lerp psyllid bugs *Glycaspis brimblecombei* [25]. [26] simulated rainfall on potato aphids *Macrosiphum euphorbiae* using a shower head, and showed that rainfall increased the amount of walking done by individuals. Here, we describe and evaluate a simple nozzle-pumped design, and use it to explore how individual and colony-level bumblebee behaviour changes in response to the presence or absence of simulated rain.

## METHODS

### Study species and animal husbandry

Three colonies of commercially reared buff-tailed bumblebees *B. terrestris* subsp. *audax* (Biobest NV, Westerlo, Belgium) housed in cardboard nest boxes with the supplied nectar reservoir removed were used, following the general husbandry and conditions described in Lawson et al. [27,28]. They were kept in a laboratory lit with UV-emitting lightbulbs on a 12 hour light:dark cycle, maintained at a temperature of approximately 21°C and relative humidity (RH) of 40%. A transparent, gated tube (300 × 15 mm) connected the nest box to a plywood flight arena (length 1140 × width 780 × height 605 mm), with green tape covering the floor and a UV-penetrable lid. Bumblebee colonies were permitted an acclimation period of two weeks before experiments began to account for transport disruption. Once sampling was complete, bumblebee colonies were euthanised by freezing.

On week days (Monday-Friday), the bees were fed 30% sucrose in the wells of artificial flowers and a PCR tray provided *ad libitum* and refreshed daily, and pollen (Sevenhills Wholefoods, Wakefield, UK) was added directly into the nest box three times a week. During experiment days, prior to experiments the availability of sucrose solution was reduced by filling fewer of the PCR tray wells, in order to encourage foraging. Alongside the PCR rack, six artificial flowers (also filled with sucrose solution) were placed within the flight arena to encourage exploratory foraging. The artificial flowers consisted of a plastic disc, with an Eppendorf tube lid containing sugar solution. To prevent nectar dilution from the rain simulator, plastic pipette ends were used to cover the artificial flower in all flight arenas (Supplementary Figure 1). In order to identify individuals, foraging bees were marked on the centre of the thorax, with different colours of non-toxic marking paint (E. H. Thorne, Rand, UK). For weekend feeding, all colonies were fed a more concentrated BIOGLUC® solution (Biobest NV, Westerlo, Belgium) *via* drip feeders. These feeders were half filled with the solution and set up on Friday afternoon as well as being given pollen and a PCR tray.

### Rain simulator design

The rain simulator consists of a flight arena (1140 mm length × 770 mm width × 305 mm height), containing a drip-irrigation system (GARDENA, Ulm, Germany). This was constructed using 3 GARDENA micro-drip nozzles angled at 45° along either side of the arena to mimic falling rain on either side of the arena (Supplementary Figure 2), connected via T-joints and micro-drip tubing (4.6 mm in diameter). The irrigation system hung using hooks and bamboo poles 180 mm high, from the walls of the flight arena. A ‘floor’ was introduced to the arena so that the bees only had access to the top 3050 mm of the space. This floor was covered with black capillary matting and reinforced with 20 × 20 mm wire mesh underneath, such that water could pass easily through into the space below. In the area below the floor, we placed two square trays 75% filled with tap water. Each tray contained one 1000 L hr^-1^ submersible water filter (Discoball, Foshan City, China) used to pump water through the system, meaning that the water within the arena was reused throughout the experiment (and changed at regular intervals). Following pilot tests, the spray from the nozzles on either side of the arena was sufficiently powerful to reach just over half way across the arena, meaning that when the pumps were operating, most of the flight space within wet area of the arena received rain from one of the nozzles. The ambient temperature and relative humidity in the arena were monitored every minute using an EasyLog USB-2 data logger (Lascar Electronics Ltd, Whiteparish, UK) to a resolution (± error) of 0.5 ± 0.55 °C and 0.5 ± 2.25 %RH, respectively. A cut-off plastic pipette dropper covered the wells of the artificial flowers to prevent nectar dilution by rainwater (figure 2). When colonies were used with the rain simulator arena, the nest box was plugged into the arena such that the bees experienced at least a day with the arena before any observations were recorded.

### Behaviour of bumblebee colonies experiencing artificial rain

Bumblebees were provided with 30% sucrose solution in approximately one-third of the wells of a PCR tray, placed out of the reach of the rain, and six randomly distributed artificial flowers. The wells were refilled after each sample to prevent food availability becoming a limiting factor and the flower positions randomised. Bumblebee colonies were exposed to 20 minutes of rainfall (where the rain machine was on for the full interval) or 20 minutes of non-rainfall (where the rain machine was completely off for the full interval), using a randomised block design. Behavioural data were collected for the final five minutes at the end of each 20-minute period, and the humidity and temperature were recorded for the first minute of this five minute block.

Before the start of every interval a new PCR tray one third filled with sugar solution was placed into the flight arena and the previous PCR tray was removed, the six artificial flowers were also emptied of sugar solution, wiped clean (to remove scent marks of previous foragers [29] and avoid microbial growth) and refilled, to avoid foraging bias. Access to the arena was barred before each interval and the number of individuals present was counted. Once the observations started, the bar was removed, letting the bees move freely between the nest and arena. Throughout the interval, the entries to and exits from the arena were counted; defined as the movement of the bumblebee’s entire body through the access point. The proportional change of active workers (those in the flight arena) during each interval was calculated by dividing the net movement (number of entries minus number of exits) by the number of bees in the arena at the start of the interval. The number of bees conducting each of four different behaviours were also recorded during these periods: flying, where the bees were in flight, not touching any surface of the flight arena; foraging, where the bees were seen extending their probosces on artificial flowers or the PCR tray; crawling, where the bees moved along the floor, sides, or roof of the flight arena; and resting, where the bees were stationary, located on the floor, roof, sides or in the corners of the flight arena. 48 data points were collected from two colonies (28 from one, and 20 from the other).

### Behaviour of individual bumblebees experiencing artificial rain

Individual bees from two colonies were provided with eight randomly distributed artificial flowers filled with 30% sucrose (Supplementary Figure 2) which were refilled and rearranged randomly after each sample. When in use, the colonies were attached to the rain simulation arena and allowed to forage normally as described above when experimental manipulations were not occurring. During the experimental period, the flight arena was cleared of all non-participating individuals before the experiment took place to avoid the confounding effects of social cues [57], so only one identifiably marked worker was present in the arena during sampling. Each worker was only used as experimental subject once, and was allowed to forage as a non-participating colony member once it had been observed. A total of 65 individuals, 32 from one colony and 33 from the other, were randomly assigned a rain or dry period in which they experienced the flight arena. At the start of an observation period, the rain simulator was switched on if a period of rainfall was selected (using a random number generator) before the marked forager was released into the flight arena.

The time that 65 individuals spent flying, foraging, resting, and crawling (as described above) was recorded, but crawling and resting were subdivided to record the height of the individual as well: ‘high’ and ‘low’ denoted whether the individual was above or below the irrigation system. Crawling was recorded to represent non-flying travel. Recording ceased when the bee attempted to return to the nest box through the gated tube, or when 20 minutes had elapsed; 10 bees remained in the arena for the entirety of this time. The total time spent in the flight arena and number of floral visits was also recorded, where a floral visit was defined as an individual landed on an artificial flower with its proboscis extended.

### Statistical analysis

All analyses were conducted using *R* 4.3.0 [30] – the analytical code and datasets are freely available at *figshare* [31]. The temperature during rain or non-rain for each of the colony behaviour observations were compared using Welch’s *t*-test, and the relationship between rain and humidity was tested with a Wilcoxon test as the data did not fit assumptions of normality.

The group behaviour data concerning the proportions of individuals conducting behaviours when rain was present or not was compositional, but data could not be transformed for a suitable compositional analysis. Instead, after checking that test assumptions were met, we conducted a principal component analysis considering the numbers of individuals conducting each of the four behaviours at the assay timepoint. The first two principal components (PC1 and PC2) were found to explain 97% of the variance, and were extracted. After log-transforming PC1 to satisfy test assumptions, we conducted a multivariate analysis of variance (MANOVA) to explore the effect of rainfall on PC1 and PC2 combined.

The data we collected for individual behaviours presented lots of observations where the individual bees did not conduct several of the behaviours (out of the six possible behaviours that could be seen during a single trip outside the nest, the mean (± SE) number of behaviours conducted was 2.66 ± 0.18), which meant that the proportional data were both highly skewed and contained a large number of zeroes. We therefore conducted a zero-augmented Dirichlet mixed effect model [32] using *zoid* 1.2.0 [33] to explore whether rainfall would impact on the behaviour of individuals, comparing model fit between a model including rain and a minimal model (running each model with four chains and 10,000 iterations). The performance of the models was compared using leave-one-out cross-validation information criterion statistics, using *loo* 2.6.0 [34,35]. To identify which behaviours were impacted, we calculated the proportion of time each individual conducted the six behaviours as if they were independent of each other, and compared how individuals experiencing rain and non-rain conditions differed in the proportion of time they uniquely conducted on each behaviour, along with the total length of time they spent in the arena, using a Mann-Whitney test (as the data were extremely non-normal), presenting approximated *p*-values where ties were present.

## RESULTS

### Effects of rain simulation on arena temperature and humidity

The foraging arena was marginally colder when the rain machine was operating (mean temperature with rain (±SE): 21.08 ± 0.05; non-rain: 21.40 ± 0.06; *t*_44.3_ = 4.03, *p* < 0.001), and humidity increased (median humidity with rain: 103.0, interquartile range 102.5 - 103.5; non-rain: 102.5, interquartile range 101.9 - 103.0; *W* = 2304, *n* = 24, 24, *p* < 0.001).

### Group

The principal component analysis suggests that rain has a big effect on the number of bees flying, demonstrated by the two ellipses in figure 1 showing virtually no overlap along the axis defined by the ‘flying’ vector and also PC2 (the lack of overlap is confirmed by the MANOVA: approximate *F*_2,45_ = 35.447, *p* < 0.001, of which PC1: *F*_1,46_ = 1.82, *p* = 0.184; PC2: *F*_1,46_ = 59.79, *p*i < 0.001). Bees were therefore much less likely to fly when it was raining (figure 2). Rain also meant bees were more likely to be very stationary (as seen with the elongated ellipse along the stationary axis in figure 1, and the difference visible in figure 2). Neither crawling nor foraging behaviour were impacted by whether rain was being experienced (figures 1, 2).

**Figure 1.**
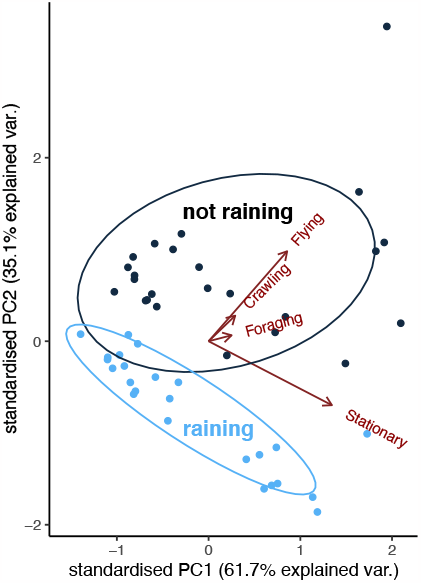
Loading plot showing contributions of the first two principal components identified as contributing to explaining the differences in colony behaviour when experiencing rainy or dry conditions.

**Figure 2.**
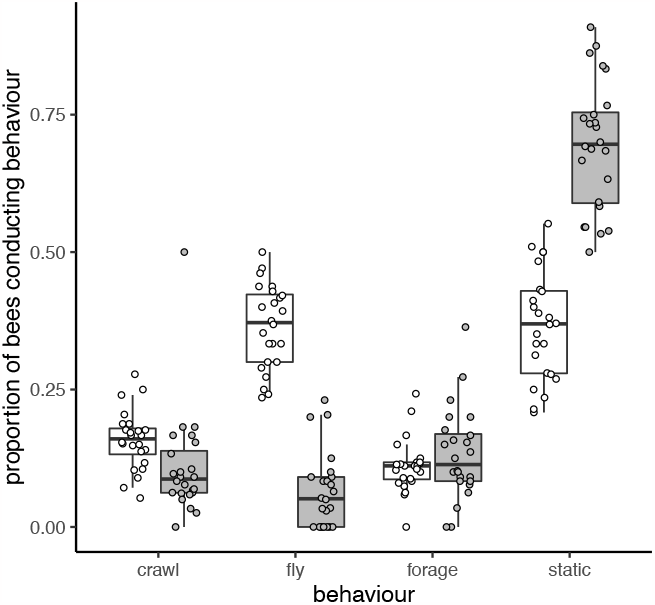
Colony response to rain, showing the proportions of the bees in the arena when there was no rain (clear bars and points) or when rain was being simulated (shaded bars and points). Boxplots show the median and interquartile values of the mean rescaled behavioural metric, and the tails show 1.5 × interquartile range.

### Individual

Models including rain as an explanatory variable performed better than the null model (figure 3, LOOIC estimate (± SE) for minimal model: 3041.4 ± 190.6; model including rain: 2980.3 ± 183.9), although we note that there is overlap in the error distributions. Bees did not differ in the length of time they spent in the arena under rain and non-rain treatments (*W* = 520, *n* = 32, 33, *p* = 0.242). They did not differ in the proportion of time they spent flying (*W* = 599, *n* = 32, 33, approximate *p* = 0.352, figure 3), crawling low (*W* = 491, *n* = 32, 33, approximate *p* = 0.616, figure 3), stationary high (*W* = 525, *n* = 32, 33, approximate *p* = 0.965, figure 3) or stationary low (*W* = 520, *n* = 32, 33, approximate *p* = 0.867, figure 3), mostly only foraged when it was not raining (*W* = 660, *n* = 32, 33, approximate *p* = 0.019, figure 3), and were more likely to crawl high in the arena when it was not raining (*W* = 701, *n* = 32, 33, approximate *p* = 0.014, figure 3).

**Figure 3.**
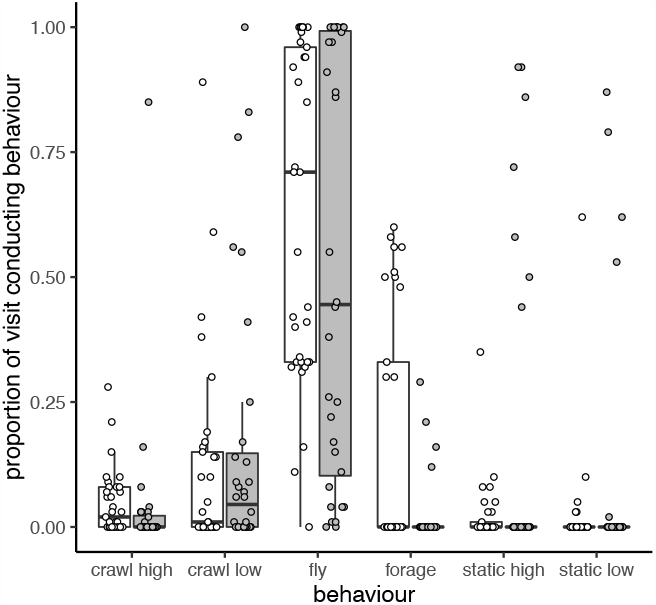
Individual bee response to rain, showing the proportions of the bees in the arena when there was no rain (clear bars and points) or when rain was being simulated (shaded bars and points). Boxplots show the median and interquartile values of the mean rescaled behavioural metric, and the tails show 1.5 × interquartile range.

## DISCUSSION

### Behavioural changes by bumblebees in response to artificial rain

The proportion of active *B. terrestris* workers decreased by ∼20% following a rain period (figure 2), supported by similar observations of bumblebees [36] and other flying insects, including Asiatic honey bees *A. cerana* [37], European honey bees *A. mellifera* [38–40] and the wasp *Vespula germanica* [41]. These findings imply that the unfavourable conditions encouraged a reduction in activity, in which some workers returned to the nest. The proportion of workers increased by ∼34% during the dry periods (figure 2), suggesting they increased their foraging effort to compensate for the reduction during the rain, echoing the activity pattern seen in *A. mellifera* workers during poor conditions [39,42].

When bees were studied individually, rain had an impact on the overall behaviour of the bees, and a greater proportion of active workers tended to forage or crawl during dry conditions (figure 3). These differences suggest variation amongst workers as some continue foraging during rain but the majority either rest until the weather improves or return to the nest altogether (figure 2). While such differences cannot be identified from this study alone, the alloethism of bumblebees, whereby larger workers forage and smaller workers perform nest duties [43], might contribute to the variation. Some species of *Bombus* bumblebees such as *B. terrestris* can forage several kilometres away from their nests [44–48], so resting during rainfall might prevent the high energetic costs of flying back to the nest [49]. *B. terrestris* and *B. impatiens* workers are hardy pollinators, showing larger flight ranges and a much wider range of plant species visited than many other pollinators [50]. Their physiology may suit them to being able to cope with poor weather conditions: this is seen in the pollinator community shift in blueberry *Vaccinium corymbosum* farmlands [13], where honeybees are the dominant species in dry weather, but *B. impatiens* tend to dominate when weather conditions are bad. Larger workers are also more efficient at foraging than smaller workers [43], suggesting that the increased forage they gather might outweigh the energetic costs of continuing to forage during the rain. Measuring the thorax width of workers making these behavioural decisions [43] would reveal whether there is a relationship between worker size and the ability of individuals to continue activity during the rain, and could help to explain the differences seen in this study. Therefore, alloethism of bumblebee workers might make them more successful pollinators than *A. mellifera* workers, which express temporal polyethism [51], during precipitation. This would also explain the relative lack of changes in behaviour shown by individual bees in response to rain in the current study – changes in deployment of different individuals could contribute to the colony’s response to precipitation.

Goulson *et al*. [43] noticed that rain extended the time of foraging trips made by *B. terrestris* workers, and concluded that this was due to sheltering in the field. However, bumblebee resting behaviour has rarely been studied in greater detail, leaving its adaptive function to be revealed. A hairy body may increase in weight when the hairs get wet, leading to greater flight costs. Optimal foraging theory predicts that the behaviour of an individual carrying an increased weight should change in response to the mass carried [52,53]. Hagen *et al*. detected that bumblebee workers with an added weight, such as a small tracking device, rested for longer than those without [46]. Larger workers can bear a greater foraging mass than smaller workers [43] so could continue foraging at a lesser cost whilst wet. Bumblebees also rest to generate the heat required for flight [50], which would take longer with a wet body due to evaporation. These mechanisms may be partially responsible for the high proportion of resting workers seen during rainfall. The choice of resting place may also be adaptive; *Anopheles* mosquitoes are more likely to initiate flight if resting on a wall or ceiling during the rain [54]. Exploring whether bumblebees actively seek rest on walls and whether they are more capable of taking off following this choice, would provide valuable insight into the adaptive resting behaviour of bumblebees.

The typical percentage of a bumblebee colony foraging during the day is around 30% [43] which is higher than was seen during this study (figure 3) and particularly more so than during rainfall (figure 4). The number of active foragers is a robust proxy measure of colony fitness [55], and these results suggest that the fitness of bumblebee colonies could be jeopardised by the conditions experienced during this study [38]. The consistently high humidity probably impacted on foraging motivation, as is seen in *A. mellifera* workers [40,56]. Bumblebee foraging trip length reduces with increasing humidity [57], suggesting that a greater proportion of workers would have been foraging if the humidity was lower and more closely associated with rain treatment. Together with an abundant nectar source a short distance from their nest [58], the conditions bumblebees experienced during this study are somewhat unreflective of those experienced in nature and likely reduced the foraging motivation. Therefore, the relationship between the proportion of foraging workers and rainfall is anticipated to be more pronounced under natural conditions, where a lower humidity is experienced when rainfall is not present. Precipitation is a multifaceted variable, and many climatic variables can influence the number of foragers [59]. Sometimes, rainfall provides little explanation for variation in activity [60] and will interact with other abiotic variables such as temperature when contributing to the activity changes seen in the field [61], highlighting the necessity for comparison between laboratory and field conditions to determine the true, direct influence of rainfall. Similarly, the behaviour of the pollinator on the flower under different conditions needs closer attention – for example, honeybees reduce foraging on pollen but maintain foraging on nectar when it is raining [62], suggesting that some species may have fine-tuned behavioural responses to immediate environmental conditions.

### Evaluating the rain simulator

Simulating rain in the lab is not going to produce something that exactly matches what would be experienced by an insect on or near the ground during a real precipitation event, and we need to consider whether the stimulus that we are presenting is biologically meaningful. Changes in atmospheric pressure, temperature, humidity and dewpoint accompany the onset of precipitation [36] but were not simulated in this study. The curcurbit beetle *Diabrotica speciosa*, the true armyworm moth *Mythimna unipuncta*, and the potato aphid *Macrosiphum euphorbiae* detect changes in atmospheric pressure and alter their mating behaviour accordingly [63]. The pressure detection capability of bumblebees has not been investigated thus far, would provide clarity on whether bumblebees reduce their activity with decreasing atmospheric pressure to avoid being out in the impending rainfall [64]. This could be achieved by adapting this rain machine into a pressure controlled flight chamber [63–65] and simulating the pressure reduction before precipitation. The effects of humidity on bumblebee activity are unclear: Morandin *et al*. [66] found no relationship between bumblebee activity and humidity, Peat and Goulson [67] found a positive correlation between foraging rate and humidity and Maurer *et al*. [57] found a negative correlation between the length of foraging trip and humidity. Furthermore, floral humidity may act as a cue for bumblebees [68,69], but environmental humidity may impact on the bees’ ability to detect and respond to this cue [70]. However, it is agreed upon that high humidity renders pollen collection impossible [67,71]. There are also conflicting conclusions on the long-term effect of humidity on colony development [72,73]. These covariates make rainfall a difficult variable to simulate. Ensuring that simulated rain is analogous to natural rain requires a particular focus on the total amount of rainfall and the rainfall intensity [74].

Rain is not a uniform entity, and naturally-falling raindrops may differ greatly in many physical properties such as size, shape, kinetic energy, and velocity, along with their spatial distribution and intensity within a precipitation event [74,75] and other environmental effects such as changes in humidity and barometric pressure. The design of a rainfall simulator needs to take account of these properties in order to present a simulation that is relevant to the real world [76,77], but there is no standardisation of simulation techniques [20,77]. It has also been noted that soil erosion studies have tended to focus on simulating extreme weather events rather than what would be experienced in ‘normal’ rain conditions [19], meaning that existing simulators may not be suitable for studying the behaviour of insects in milder precipitation events. The small, similarly-sized droplets typically produced by rain simulators produce only low kinetic energy which is unreflective of natural rain [74,78], from which it can be concluded that bumblebees in this study experienced very light rain.

A consistent average relative humidity of over 100 % is rarely experienced in nature, yet the bumblebees in this experiment were able to continue their usual behaviours, contrary to the activity by *A. mellifera* workers [40,56]. Future use of the rain machine should endeavour to reduce this humidity to a value reflective of natural conditions. This is possible by using a solid floor with drainage so rainwater is not retained by capillary matting (figure 1*b*). Utilising the arena design by Lawson *et al*. [27], implementing a fan to disperse olfactory cues, would disperse some of the moisture, but could bias droplet dispersion. Silica gel petri dishes underneath the arena floor [79] would be preferable to reduce humidity if droplet dispersion is important to the study [74]. To accurately maintain humidity, a dry air stream (such as used in [80]) may also be used in conjunction with the rain machine to more accurately represent the conditions associated with precipitation.

If the behaviour of an animal is directly influenced by what sort of rainfall it is experiencing, then care would need to be taken in experiments to correctly manipulate the properties of the rain that are relevant to the animal. For behavioural studies of captive small animals, it is likely that researchers will wish to conduct their observations either in the field or within a relatively small lab space. Small (and relatively portable) simulators have been designed and used (e.g. [20,81–83]), but may be limited in their ability to simulate some physical properties of natural raindrops such as fall velocity (that requires the drop to fall a minimum distance first) [20]. We would therefore recommend that either care is taken to identify the physical properties of rainfall that are relevant to an experimental species, or that an attempt is made to measure [75,84] and standardise the simulated rainfall, acknowledging that there is likely to be a trade-off between the size of the simulation equipment and the authenticity of the rain experienced. Given that climate change is likely to lead to higher or more intense levels of rainfall [2–5], we need identify and explore all the possible effects of these changes on pollination services [7], and targeted exploration under laboratory conditions may offer us a fast and reliable means of picking apart some of these effects.

### Data availability

The dataset and *R* code used are freely available on *figshare* [31].

## Author Contributions

Conceptualization: LAR EJ DAL SAR Data curation: SAR

Formal analysis: SAR

Investigation: LAR EJ

Methodology: LAR EJ DAL SAR

Supervision: DAL SAR

Visualization: SAR

Writing – original draft: LAR EJ SAR

Writing – review & editing: LAR EJ DAL SAR

## SUPPLEMENTARY FIGURES

**Supplementary Figure 1.**
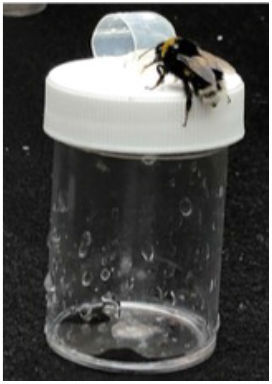
Sheltered artificial flower, showing cap placed over feeding well in order to avoid dilution of the sucrose solution reward.

**Supplementary Figure 2.**
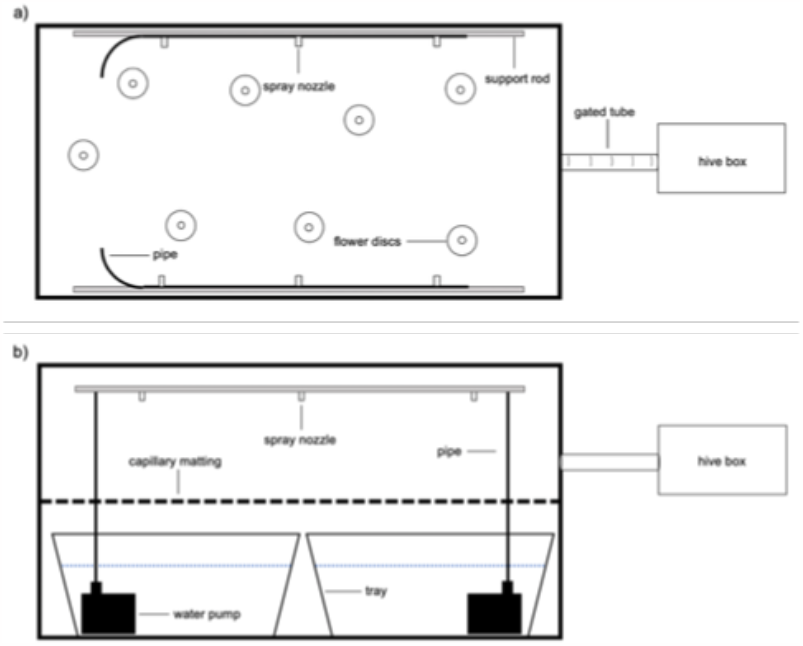
Sketches of the rain simulator. a) A top-down view of the experimental set up of a rain simulator within a bumblebee flight arena, using a modified irrigation system of spray nozzles angled at 45°. b) A side view of the same setup, showing the closed-loop water system where the water is pumped into the flight arena, which falls back into the trays *via* capillary matting supported by sturdy wire mesh to create a stable floor. Note that the flower arrangement in the arena is sketched for the individual-level experiments (where there were eight flowers present), rather than the colony-level experiments (where there were six flowers and an additional PCR tray).

